# Histo-anatomical atlas and thermal tolerance of *Garra rufa*: A novel small teleost model adaptable to human body temperature

**DOI:** 10.64898/2026.02.27.708595

**Authors:** Tetsuo Kon, Koto Kon-Nanjo, Shuma Nihei, Liqing Zang, Oleg Simakov, Yasuhito Shimada

**Affiliations:** Department of Neurosciences and Developmental Biology, University of Vienna, Djerassiplatz 1, Vienna, 1030, Austria; Department of Integrative Pharmacology, Mie University Graduate School of Medicine, 2-174 Edobashi, Tsu, Mie 5148507, Japan; Graduate School of Regional Innovation Studies, Mie University, 1377 Kurimamachiya-cho, Tsu, Mie 514-8507, Japan; Mie University Zebrafish Research Center, 2-174 Edobashi, Tsu, Mie 5148507, Japan

**Keywords:** Doctor fish, *Garra rufa*, high-temperature tolerance, histo-anatomy, model organism, red garra, zebrafish

## Abstract

*Garra rufa*, commonly known as the doctor fish, is a small freshwater cyprinid notable for its exceptional tolerance to high temperatures, surviving even at around the human body temperature of 37 °C, and has emerging potential as a novel laboratory model for human cancer xenotransplantation and infectious disease research. To establish a foundation for its experimental use, we conducted comprehensive anatomical and histological analyses across major organ systems. The overall body organization and tissue architecture of *G. rufa* are broadly similar to those of zebrafish (*Danio rerio*), indicating a conserved cyprinid body plan. However, several organ systems in *G. rufa* exhibited species-specific differences compared with zebrafish, including a well-developed adhesive disc around the oral region, a long and coiled intestine, and a distinct dark pigmentation of the peritoneum. These species-specific traits may reflect ecological and behavioral adaptations of *G. rufa*, including benthic scraping in warm, flowing habitats. Physiological assays confirmed that *G. rufa* maintains high survival rates and normal swimming activity at 37 °C, whereas zebrafish exhibit significant mortality and reduced locomotion under the same conditions. Collectively, this work provides a comprehensive histo-anatomical atlas of *G. rufa*, highlighting its unique morphological specializations while establishing an essential reference for the development of this species as a novel experimental fish model.

## Introduction

*Garra rufa* (Heckel, 1843), also known as the doctor fish or red garra, is a small freshwater cyprinid mainly found in rivers, hot springs, and thermal pools in the Middle East^1^. This fish has been used in ichthyotherapy, a treatment in which the fish feed on keratinized skin from human hands and feet, providing a natural exfoliation^2,3^. *G. rufa* exhibits remarkable tolerance to high temperatures, as demonstrated by its habitat in thermal waters and its application in ichthyotherapy, where the fish come into direct contact with human skin.

Small teleost fish, such as zebrafish and medaka, are typically only a few centimeters in length and have been widely used as vertebrate model systems in various fields of life sciences, including developmental biology^4–6^ and drug discovery^7,8^. The advantages of small teleost fish models include a rapid life cycle, high fecundity, compact housing requirements, suitability for high-throughput drug screening, and compatibility with animal welfare considerations. In recent years, alongside conventional models, several new small fish models with unique biological features have been developed, including the goldfish (*Carassius auratus*) for whole genome duplication research^9–11^, the African turquoise killifish (*Nothobranchius furzeri*) for aging research^12,13^, and the threespine stickleback (*Gasterosteus aculeatus*) for environmental adaptation research^14,15^. Building on the extensive foundation of zebrafish and medaka research, studies utilizing these emerging teleost models are rapidly advancing.

Maintaining body temperature within an appropriate range is crucial for proper biochemical reactions in cells and tissues^16,17^. In particular, tissues with high metabolic activity, such as the liver and visceral fat, are significantly affected by changes in body or ambient temperature^18–20^. Therefore, the mechanisms by which *G. rufa* can adapt to high temperatures, conditions that are harsh for other teleost fish including zebrafish (optimal at ∼28°C), are of great interest from the perspective of metabolic system evolution. Furthermore, experiments at the human body temperature of 37°C are critical for modeling human physiology when using small teleost models for studies on metabolic diseases, drug metabolism, human cancer xenografting, and human infectious diseases. For example, in zebrafish models for human cancer transplantation aimed at drug screening, the physiological temperature discrepancy between the host (zebrafish) and the human cancer cells poses a challenge for optimal tumor proliferation^21,22^.

With the tolerance to temperatures of 37°C or higher and the recently established chromosome-level genome assembly^23^, *G. rufa* holds great potential as a novel biomedical model complementing zebrafish^24^. In addition, the molecular mechanisms underlying the high-temperature tolerance of *G. rufa* itself may provide significant insights into the environmental adaptation strategies of vertebrates. However, detailed histo-anatomical analyses essential for developing *G. rufa* as a laboratory model, as well as direct comparisons of its survival and behavioral responses to 37°C with those of zebrafish, have not yet been systematically addressed. Here, we performed comprehensive histo-anatomical analyses of *G. rufa* and compared its survival and behavioral responses to high temperature with those of zebrafish.

## Results

### Gross anatomical features

*G. rufa* exhibits a streamlined body shape typical of cyprinids (Fig. 1a), similar to zebrafish (Fig. 1b). The fish displays a grayish-brown body coloration, which likely enhances camouflage within its natural riverbed environment. *G. rufa* possesses dorsal, pectoral, pelvic, anal, and caudal fins that facilitate locomotion (Fig. 1b). The dorsal fin is located along the midline, while the pectoral fins are positioned laterally, posterior to the operculum (gill cover). The pelvic fins are situated ventrally near the mid-body, and the anal fin is located on the ventral side, just anterior to the caudal fin. Whole-body micro-computed tomography (µCT) imaging revealed that these fins are supported by skeletal elements (Fig. 1c, S1). Magnetic resonance imaging (MRI) visualized the spatial relationships among the internal organs, such as intestine and swim bladder, as well as the presence of air-filled lumens (Fig. 1d, S2). A lateral line system extends along the flanks, allowing the fish to detect water currents and vibrations, thereby aiding in navigation and prey detection (Fig. 1b). The eyes are positioned laterally, providing *G. rufa* with a wide field of peripheral vision, presumably for spotting both predators and prey (Fig. 1b). Notably, *G. rufa* features an inferior mouth, with the lower lip modified into an adhesive disc (Fig. 1b). This adhesive disc enables *G. rufa* to attach firmly to underwater surfaces and scrape biofilms or algae for feeding in flowing water environments^25^.

**Figure 1.**
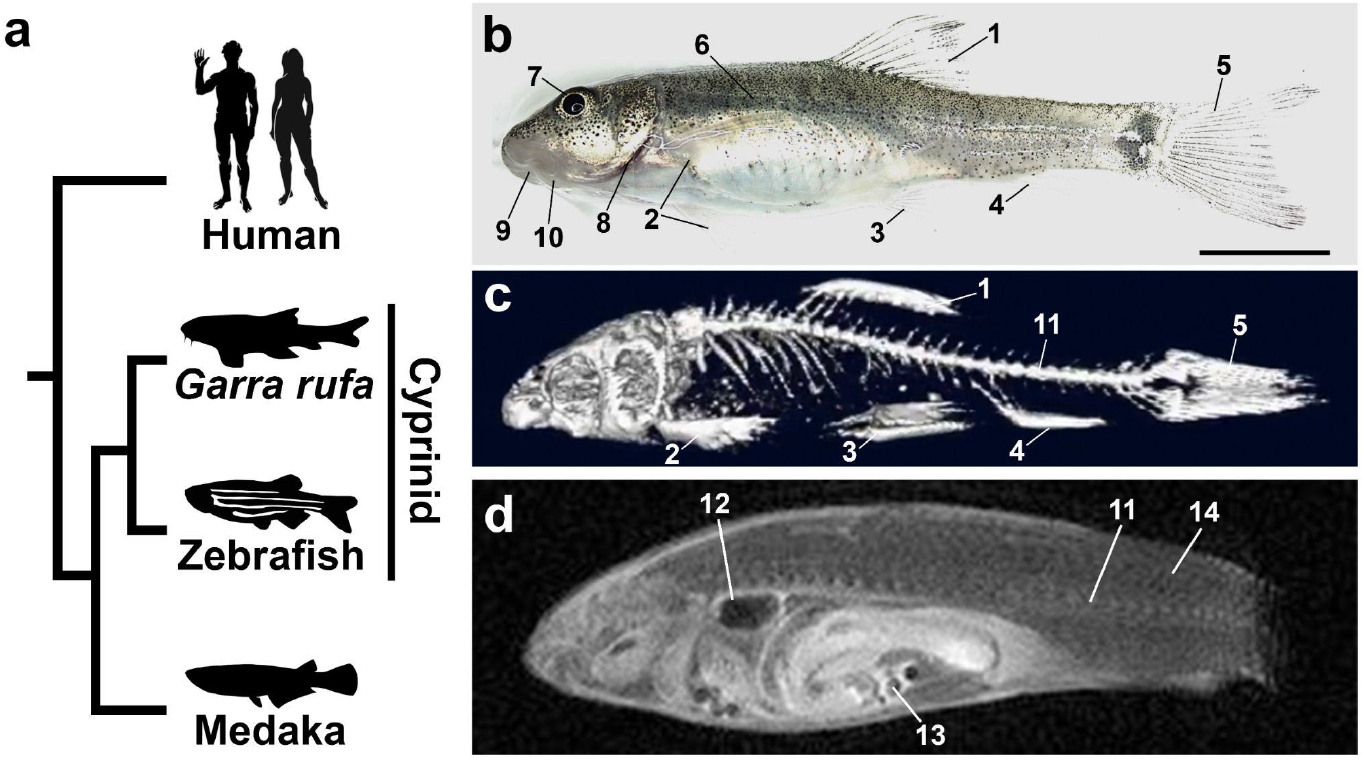
Gross anatomy of *G. rufa*. (**a**) Phylogenetic position of *G. rufa*. Animal silhouettes were obtained from the PhyloPic (https://www.phylopic.org/). (**b**) External anatomy of *G. rufa*. The scale bar represents 5 mm. (**c**) Whole-body µCT image. (**d**) T1-weighted MRI image. 1, dorsal fin; 2, pectoral fin; 3, pelvic fin; 4, anal fin; 5, caudal fin; 6, lateral line; 7, eye; 8, operculum; 9, mouth opening; 10, adhesive disc; 11, vertebral column (spine); 12, swim bladder; 13, intestine; 14, muscle tissue.

### Histo-anatomical findings

To characterize the internal anatomy of *G. rufa* in detail, we conducted gross anatomical observations of the visceral organs (Fig. S3a–l). In addition, tissue sections were prepared and stained with hematoxylin and eosin (H&E) to examine the histological architecture of each organ or tissue (Fig. S4a–e). We also utilized publicly available zebrafish virtual slides^26^ to compare conserved structures between *G. rufa* and zebrafish, and to identify morphological features unique to *G. rufa* (Fig. S5a–w, Table S1).

### Digestive tract

#### Mouth

The lower lip of *G. rufa* is modified to form an adhesive disc (Fig. 2a–c). This disc is circular and features a smooth, shallow central depression, which functions as an effective structure for adhesion. This adaptation facilitates adherence to rocks and pebbles in flowing water, thereby supporting the benthic lifestyle of this fish^25^. The adhesive disc is covered by stratified squamous epithelium (Fig. 2c). In contrast, zebrafish do not exhibit such specialization of the lower lip for adhesion to surrounding substrates (Fig. S5a), which is consistent with their pelagic, free-swimming behavior rather than a benthic lifestyle^27^.

**Figure 2.**
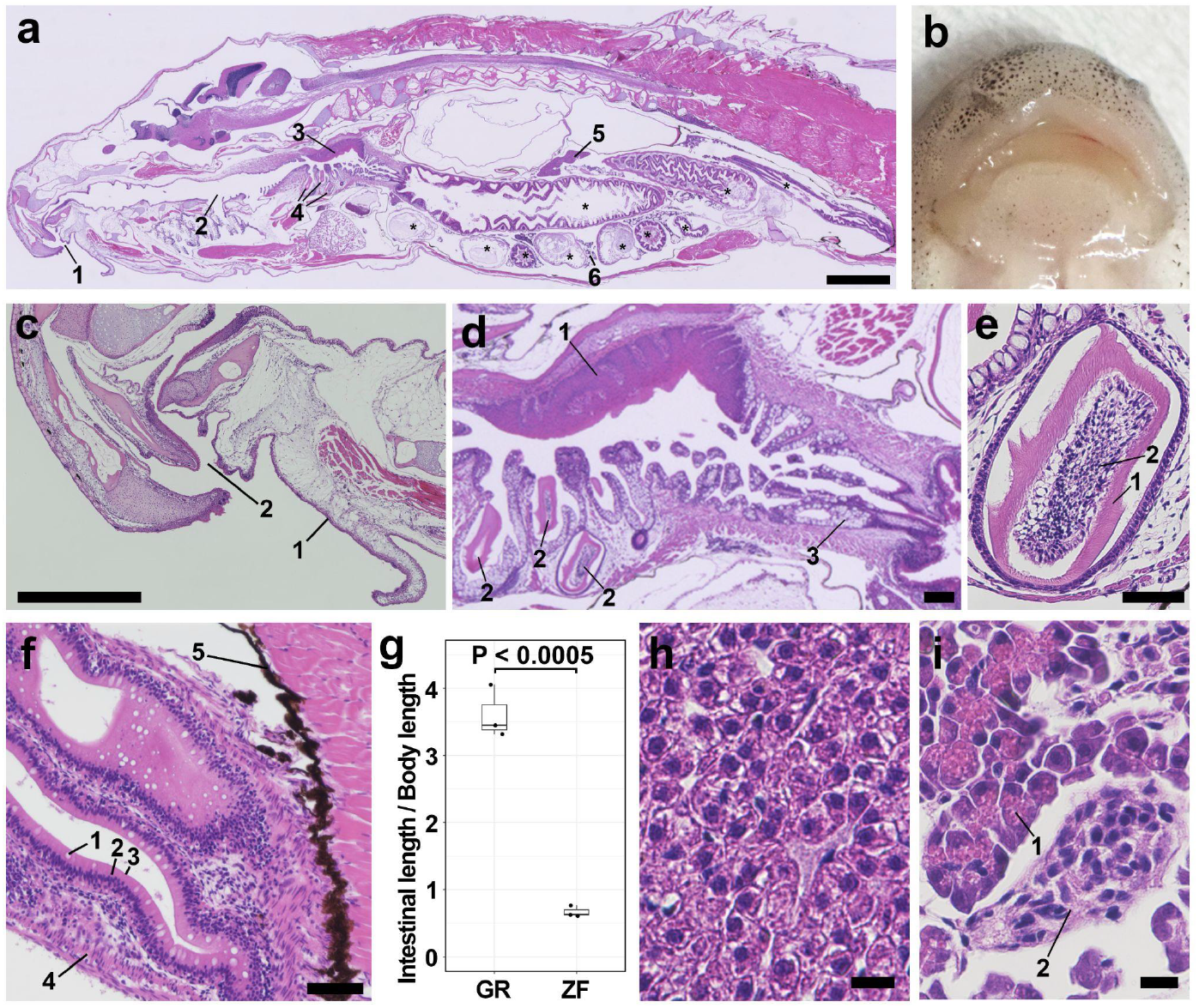
Digestive tract and glands associated with the digestive tract. (**a**) Low-magnification image showing the overall arrangement of the digestive tract. 1, mouth opening; 2, oropharynx; 3, pharyngeal pad; 4, pharyngeal tooth; 5, liver; 6, pancreas in the mesentery. Asterisks indicate the intestine. Scale bar: 1 mm. (**b**) Gross morphology of the mouth and the adhesive disc. (**c**) Sagittal section of the mouth region showing the adhesive disc. 1, adhesive disc; 2, mouth opening. Scale bar: 500 µm. (**d**) Pharynx and esophagus. 1, pharyngeal pad; 2, pharyngeal tooth; 3, goblet cells in the esophagus. Scale bar: 100 μm. (**e**) Pharyngeal tooth. 1, dentine layer; 2, pulp cavity. Scale bar: 50 μm. (**f**) Peritoneum and intestine. 1, goblet cells; 2, enterocytes; 3, brush border; 4, smooth muscle layer; 5, peritoneum. Scale bar: 50 μm. (**g**) Comparison of intestinal length between *G. rufa* (GR) and zebrafish (ZF). (**h**) Liver. Hepatocytes are densely arranged without a clear lobular or cord-like organization. Scale bar: 10 μm. (**i**) Pancreas. 1, acinar cells; 2, islet of Langerhans. Scale bar: 10 μm.

#### Oropharynx

The oropharynx constitutes the transitional segment between the oral cavity and the esophagus. Histologically, the oropharyngeal epithelium contains numerous goblet cells (Fig. 2d). A distinct pharyngeal pad is present on the dorsal wall at the posterior part of the oropharynx. The pharyngeal pad consists of a thickened mucosa with stratified squamous epithelium (Fig. 2d). Pharyngeal teeth are located ventral to the pharyngeal pad (Fig. 2d). The pharyngeal teeth exhibit a dentine layer and a central pulp cavity (Fig. 2e). The pharyngeal teeth lack roots. These teeth presumably work together with the pharyngeal pad to grind ingested food^28^. Zebrafish also possess pharyngeal teeth and a pharyngeal pad (Fig. S5b,c).

#### Esophagus

The esophagus extends from the oropharynx to the anterior segment of the intestine (Fig. 2d). The esophagus shows prominent mucosal folds, a feature characteristic of the highly distensible teleost esophagus, which permits the swallowing of substantial quantities of food and water^28^. In zebrafish, the esophagus similarly exhibits pronounced mucosal folds (Fig. S5b). The esophageal wall is organized into layers from the lumen outward, comprising the mucosa, the muscle layer, and the serosa (Fig. 2d). The esophageal epithelium is composed of epithelial cells and numerous goblet cells.

#### Abdominal cavity

The abdominal cavity contains organs such as the intestine, liver, pancreas, and spleen (Fig. S3a). Its surface is lined by the peritoneum. Notably, in *G. rufa*, extensive black pigmentation is observed on the peritoneal surface (Fig. S3b). Black pigmentation of the peritoneum has also been reported in other *Garra* species^29^. In contrast, the peritoneum of zebrafish is pale and lacks such extensive black pigmentation (Fig. S3l). Microscopically, the peritoneum of *G. rufa* shows extensive melanin deposition, which is consistent with macroscopic observation (Fig. 2f).

#### Intestine

The intestine serves as the main organ responsible for digesting food and absorbing nutrients^30^. Intestinal morphology has been reported to vary greatly among teleost species depending on their dietary habits^31^. The intestine is continuous with the esophagus and is located ventral to the swim bladder (Fig. 2a). Neither *G. rufa* nor zebrafish exhibit an anatomically distinct stomach (Fig. 2a), which is consistent with the fact that cyprinids are well known to be agastric teleosts^32^. Macroscopically, the intestine of *G. rufa* is highly coiled (Fig. S3a,c). This contrasts with the macroscopically simple and largely straight intestine of zebrafish, which possesses only two major bends defining the conventional anterior, mid, and posterior segments^28,33^. When comparing the intestinal length normalized by body length between *G. rufa* and zebrafish, the average normalized intestinal length of *G. rufa* was 5.4-fold greater than that of zebrafish (95% CI: 3.6–7.1; Student’s t-test, *p* < 0.0005; Fig. 2g). Adipose deposition in the mesentery is also markedly prominent in *G. rufa* (Fig. S3c).

The intestinal wall contains the mucosa and the muscle layer (Fig. 2f). The mucosa comprises an epithelial layer and the lamina propria beneath it. In the epithelial layer of the mucosa, the columnar-shaped absorptive enterocytes are the most abundant cell type (Fig. 2f). Enterocytes possess brush borders, which likely facilitate nutrient absorption. Goblet cells are scattered between the enterocytes (Fig. 2f). The lamina propria consists of loose connective tissue. The muscle layer consists of two layers of smooth muscle fibers, an inner circular layer and an outer longitudinal layer, which together support intestinal movements.

### Glands associated with the digestive tract

#### Liver

The liver is a vital organ with broad physiological functions, encompassing metabolism, detoxification, and the synthesis of essential proteins^34^. Macroscopically, the liver forms flat, elongated, and branched lobes that overlay the intestine within the body cavity (Fig. S3c). Microscopically, *G. rufa* hepatocytes lack a distinct lobular or cord-like arrangement, and the portal areas are indistinct (Fig. 2h), in contrast to the well-organized lobular architecture characteristic of mammals^35,36^. The hepatocytes exhibit an irregular polygonal morphology with rounded nuclei. The liver parenchyma is compact and contains minimal connective tissue. The liver of the zebrafish displays a histological organization similar to that of *G. rufa* (Fig. S5e).

#### Pancreas

In both teleosts and mammals, the pancreas secretes digestive enzymes into the intestinal tract to facilitate food digestion and also functions as an endocrine organ by producing multiple hormones^35,36^. Unlike mammals, which possess a compact pancreatic organ^36,37^, the pancreatic tissue in *G. rufa* is diffuse, existing as scattered islands embedded within the mesenteric adipose tissue, a feature typical of other teleosts^38^. Histologically, the pancreas consists of both endocrine and exocrine components (Fig. 2i). The acinar structures of the exocrine component contain acinar cells with basophilic cytoplasm and eosinophilic zymogen granules (Fig. 2i). Islets of Langerhans are interspersed among the exocrine components (Fig. 2i).

### Cardiovascular system and blood

#### Heart

The vertebrate heart is a continuously beating organ responsible for circulating blood throughout the body^39^. In *G. rufa*, the heart is located ventral to the esophagus and posterior to the gills (Fig. 3a, S4). Macroscopically, the heart of *G. rufa* consists of four compartments arranged in series: the sinus venosus, atrium, ventricle, and bulbus arteriosus (Fig. 3a). This structure contrasts with the four-chambered hearts of mammals^35,36,39^. The ventricular cavity contains well-developed trabeculae, and the cardiac wall is composed of myocardium formed by cardiac muscle cells (Fig. 3a). Zebrafish exhibit a cardiac structure essentially similar to that of *G. rufa* (Fig. S5g), reflecting the conserved organization of the teleost heart.

**Figure 3.**
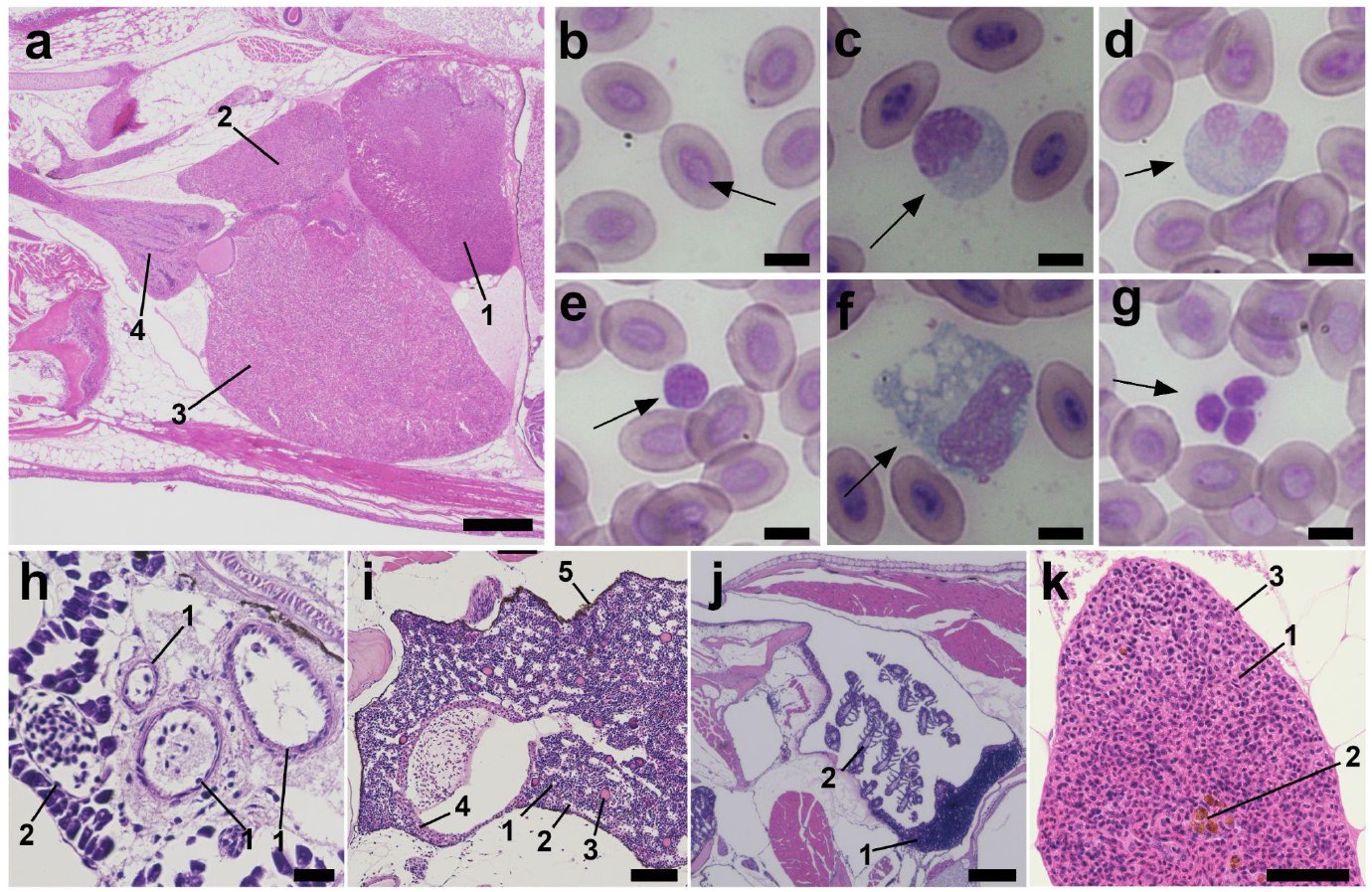
Heart, peripheral blood and immune organs. (**a**) Heart. Scale bar: 200 μm. 1, sinus venosus; 2, atrium; 3, ventricle; 4, bulbus arteriosus. (**b**) Erythrocyte (arrow). Scale bar: 10 µm. (**c**) Neutrophil (arrow). Scale bar: 10 µm. (**d**) Eosinophil (arrow). Scale bar: 10 µm. (**e**) Lymphocyte (arrow). Scale bar: 10 µm. (**f**) Macrophage (arrow). Scale bar: 10 µm. (**g**) Thrombocyte (arrow). Scale bar: 10 µm. (**h**) Arteriole and venule. 1, arteriole; 2, venule. Scale bar: 20 μm. (**i**) Pronephros. 1, lymphatic tissue; 2, blood sinus; 3, thyroid follicles; 4, adrenal tissue. Scale bar: 100 μm. (**j**) Thymus. 1, thymus; 2, gill. Scale bar: 200 μm. (**k**) Spleen. 1, splenic parenchyma; 2, melanomacrophage center; 3, splenic capsule. Scale bar: 50 μm.

#### Peripheral blood

To evaluate the cellular composition of peripheral blood, blood smears were prepared and stained with Wright-Giemsa (Fig. 3b–g). The most abundant cell type observed was the nucleated erythrocytes (Fig. 3b), a characteristic feature shared by many teleosts, including zebrafish^40^. This contrasts with the enucleated red blood cells found in mammals. In addition to erythrocytes, immune cells such as neutrophils (Fig. 3c), eosinophils (Fig. 3d), lymphocytes (Fig. 3e), and macrophages (Fig. 3f) were identified. Nucleated thrombocytes were also present (Fig. 3g).

#### Blood vessels

As in other vertebrates, the vascular system of *G. rufa* comprises three main types of blood vessels: arteries, veins, and capillaries. The arterial system is characterized by endothelial vessels surrounded by smooth muscle layers of variable thickness (Fig. 3h), encompassing both arteries and arterioles. The venous system consists of vessels lined by endothelium with relatively thinner smooth muscle layers, including veins and venules. Veins typically run parallel to arteries (Fig. 3h). Capillaries have the thinnest vessel walls and are distributed throughout the body, including the gills and kidneys, allowing efficient transport of oxygen and nutrients.

### Immune organs

#### Pronephros

The pronephros (head kidney) is a paired lymphohematopoietic organ located in the anterior dorsal body cavity, just posterior to the gills (Fig. 3i,S4). In teleosts, the pronephros primarily functions as a hematopoietic tissue^41^. Histologically, it contains abundant lymphocytes and granulocytes, while renal tubules and glomeruli are largely absent (Fig. 3i). The pronephros is surrounded by a thin capsule with melanin deposition (Fig. 3i). The microscopic organization of the *G. rufa* pronephros is closely similar to that of the zebrafish pronephros (Fig. S5h).

#### Thymus

The thymus consists of a pair of lymphoid organs positioned along the dorsal wall of each branchial cavity (Fig. 3j). This arrangement contrasts with that of the mammalian thymus, whose precursor cells migrate from the pharyngeal region into the thorax and fuse at the midline to form a single organ^42^. The cortex and medulla are not clearly separated in the thymus of *G. rufa*. Histologically, it consists of densely packed lymphocytes, consistent with its role as a primary lymphoid organ involved in lymphocyte development. Zebrafish possess anatomically and histologically similar paired thymi (Fig. S5i)

#### Spleen

The spleen is a compact, dark-red organ located in the abdominal cavity (Fig. S3e). Histologically, the spleen is populated by numerous immune cells, consistent with its role as a major immune organ (Fig. 3k). Melanomacrophage centers are present within the parenchyma, and these structures are thought to be associated with immune responses^43,44^. Melanomacrophage centers appear more abundant in *G. rufa* than in zebrafish. (Fig. 3k, S5j).

### Respiratory system

#### Gills

The gills serve primarily as respiratory organs, but they also regulate salt and water exchange and contribute to nitrogenous waste excretion in teleosts^38,45^. The gills are symmetrically positioned in the anterior region of the oropharynx (Fig. S4). The gill of *G. rufa* is composed of gill arches, gill rakers, and primary lamellae (gill filaments) (Fig. 4a, S3i). Each primary lamella is supported by bony tissues (Fig. 4a). The gill rakers are located on the medial side of the gill arches, and exhibit a serrated morphology (Fig. 4a), facilitating food filtration^28^. Microscopically, secondary lamellae arise from the primary lamellae, and the capillaries within these secondary lamellae serve as the sites of gas exchange, which is supported by unidirectional water flow across the gills (Fig. 4a).

**Figure 4.**
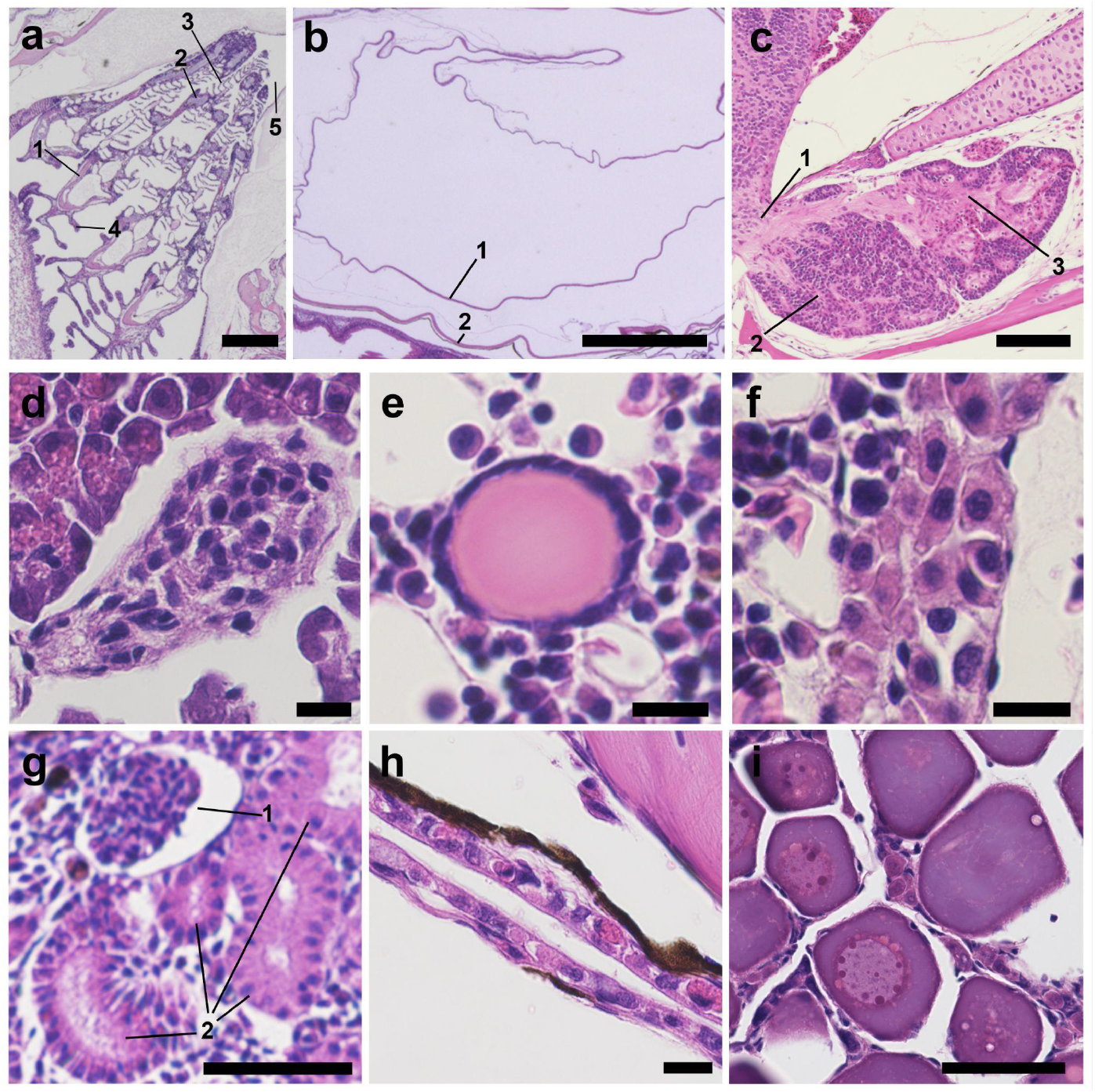
Respiratory, endocrine, urinary, and reproductive systems. (**a**) Gills. Scale bar: 100 μm. 1, gill arch; 2, primary lamella (gill filament); 3, secondary lamella; 4, gill raker; 5, gill cavity. (**b**) Swim bladder. 1, mucus membrane; 2, outer fibrous layer. Scale bar: 500 μm. (**c**) Pituitary gland. 1, pituitary stalk; 2, anterior pituitary; 3, posterior pituitary. Scale bar: 100 μm. (**d**) Islet of Langerhans. Scale bar: 50 μm (**e**) Thyroid follicle. Scale bar: 10 μm (**f**) Adrenal tissue. Scale bar: 10 μm (**g**) Kidney. 1, glomerulus; 2, renal tubules. Scale bar: 50 μm. (**h**) Ureter. Scale bar: 10 μm. (**i**) Ovary. Scale bar: 50 μm.

#### Swim bladder

The swim bladder is a gas-filled organ and located in the dorsal region of the body cavity (Fig. S2,S4). The swim bladder is divided into anterior and posterior chambers (Fig. S3h,S4). While the primary function of the swim bladder is buoyancy regulation, it can also function as an accessory respiratory organ that provides oxygen to the body under hypoxic conditions^46^. Multiple lines of evidence from developmental biology and gene expression studies strongly suggest that it is homologous to the mammalian lung^47^. Microscopically, the wall of the swim bladder comprises inner mucosal membrane and an outer fibrous layer (Fig. 4b).

### Endocrine system

#### Pituitary gland

The pituitary gland is a key endocrine tissue that secretes several types of pituitary hormones^48^. The pituitary gland is located in the pituitary fossa at the cranial base (Fig. 4c). The pituitary contains two embryologically distinct components: the anterior pituitary (adenohypophysis) and the posterior pituitary (neurohypophysis). Histologically, the anterior pituitary is composed of densely packed glandular cells, whereas the posterior pituitary exhibits a neural component continuous with the pituitary stalk (Fig. 4c).

#### Islet of Langerhans

The islets of Langerhans in *G. rufa* are found within the pancreatic tissue (Fig. 4d). These endocrine regions are embedded within the exocrine pancreas and are responsible for hormone production. In vertebrates, the endocrine pancreas is primarily composed of three major hormone-secreting cell types including α-cells that produce glucagon, β-cells that produce insulin, and δ-cells that produce somatostatin, together with several minor endocrine cell populations^28,34,36^. In the islets of Langerhans of both *G. rufa* and zebrafish, the endocrine cells are small to medium-sized polygonal cells with round nuclei and slightly eosinophilic cytoplasm (Fig. 4d, S5f). Subpopulations of endocrine cells are not readily distinguishable using the H&E staining.

#### Thyroid

The thyroid gland of teleosts releases thyroid hormones, which stimulate many metabolic processes^38^. The thyroid of *G. rufa* is not encapsulated but consists of scattered follicles primarily located throughout the connective tissue of the pharyngeal area and within the parenchyma of the pronephros (Fig. 4e), whereas mammals exhibit clustered follicles^35,39^. These follicles are lined with cuboidal epithelial cells and contain colloid material in the lumen (Fig. 4e).

#### Adrenal tissue

*G. rufa* does not possess a discrete adrenal gland as seen in mammals^36^. Instead, adrenal cells are scattered in association with major blood vessels and the pronephric region (Fig. 4f). These adrenal cells are presumed to consist of interrenal cells, which are considered homologous to the mammalian adrenal cortex, and chromaffin cells, which correspond to the adrenal medulla^28,34,38^. However, these two cell populations are not readily distinguishable by H&E staining.

### Urinary system

#### Kidney

In vertebrates, the kidney develops through three successive stages: the pronephros, mesonephros, and metanephros^49^. Teleost fish retain both a functional pronephros and a mesonephros in adults^28^. The pronephros (head kidney) later develops into a lymphoid organ, whereas the mesonephros (trunk kidney) remains the excretory kidney. In contrast, mammals retain only the metanephros as the permanent kidney^49^. In *G. rufa*, the kidney is located in the retroperitoneal space (Fig. S3j, S4). Histologically, the kidney contains glomeruli and renal tubules (Fig. 4g). Overall, the renal histology of *G. rufa* is similar to that of zebrafish (Fig. S5h, o).

#### Ureter

The kidneys drain directly to the ureter, which forms a single tubular structure (Fig. 4h). The ureter courses along the dorsal-most aspect of the body cavity. Zebrafish exhibit a similar anatomical trajectory and histological organization of the ureter (Fig. S5p).

### Reproductive system

#### Female gonad

The ovary is a paired organ occupying much of the abdominal cavity (Fig. S4). Histologically, the ovary is composed of oocytes at various stages of development (Fig. 4i). This is consistent with the pattern seen in many teleosts, including cyprinids, whose ovaries typically contain oocytes at multiple developmental stages simultaneously, reflecting asynchronous oogenesis^50,51^. Similarly, the ovary of zebrafish shows oocytes in multiple developmental stages (Fig. S5q).

### Nervous system

#### Brain

The brain of *G. rufa*, like that of other teleosts, is organized into major regions including the telencephalon, diencephalon, mesencephalon, metencephalon, and medulla oblongata (Fig. 5a, S3k). The telencephalon, located at the most anterior part of the central nervous system, includes the olfactory bulbs and the cerebral hemispheres. The diencephalon is located posterior to the telencephalon. The ventral region of the diencephalon is occupied by the hypothalamus (Fig. 5a). The mesencephalon is located posterior to the diencephalon. The mesencephalon contains the optic tectum (homologous to the superior colliculus in mammals) in its dorsal region for receiving and integrating visual inputs from the optic nerve^52^. The cerebellum (part of the metencephalon) is located posterior to the optic tectum. The medulla oblongata is located at the most posterior part of the brain and is involved in vital autonomic functions such as respiration and circulation in vertebrates^53^. The overall organization of the zebrafish brain is essentially similar, comprising the same major regions (Fig. S5r).

**Figure 5.**
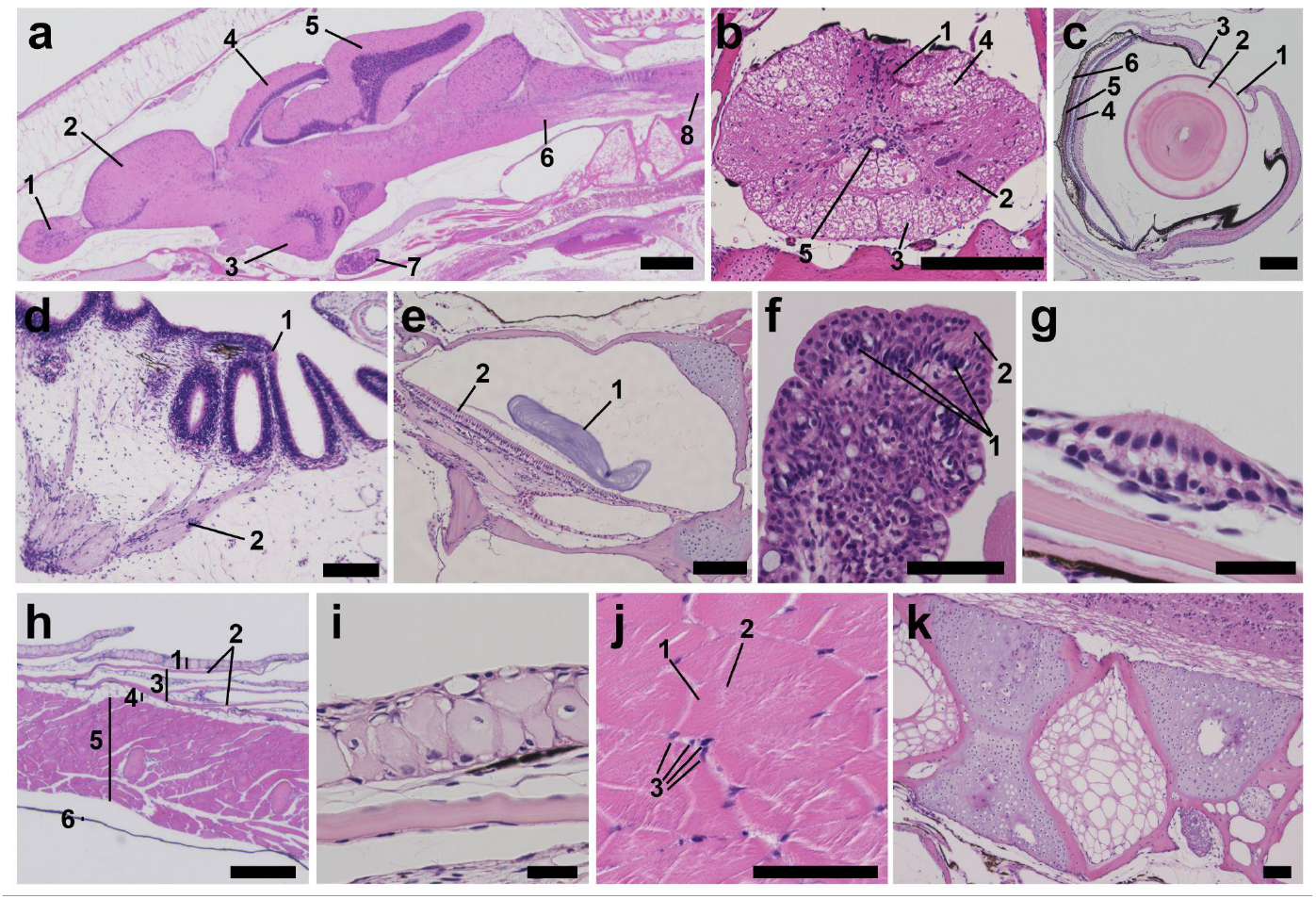
Nervous system, sensory system, skin, bone, and cartilage. (**a**) Brain and spinal cord. 1, olfactory bulb; 2, telencephalon; 3, hypothalamus; 4, optic tectum; 5, cerebellum; 6, medulla oblongata; 7, pituitary gland; 8, spinal cord. Scale bar: 500 µm. (**b**) Transverse section of the spinal cord. 1, dorsal horn; 2, ventral horn; 3, ventral funiculus; 4, lateral funiculus; 5, central canal. Scale bar: 200 µm. (**c**) Eye. 1, cornea; 2, lens; 3, iris; 4, retina; 5, retinal pigment epithelium; 6, choroid. Scale bar: 200 µm. (**d**) Olfactory sac. 1, ciliated columnar epithelial cells; 2, nerve. Scale bar: 100 µm. (**e**) Inner ear. 1, otolith; 2, sensory epithelium. Scale bar: 100 µm. (**f**) Taste buds in the pharyngeal epithelium. 1, sensory cells; 2, microvilli. Scale bar: 50 µm. (**g**) Hair cells of the lateral line. Scale bar: 20 µm. (**h**) Body wall. 1, epidermis; 2, scale; 3, dermis; 4, subcutaneous tissue; 5, skeletal muscle; 6, peritoneum. Scale bar: 200 µm. (**i**) Epidermis. Scale bar: 20 µm. (**j**) Skeletal muscle. 1, myofiber; 2, myofibril; 3, myonuclei. Scale bar: 50 µm. (**k**) Vertebra. Scale bar: 50 µm.

#### Spinal cord

The spinal cord of *G. rufa* originates from the medulla oblongata and extends caudally, running within the vertebral column (Fig. 5a). Within the spinal cord, the gray matter, which is composed mainly of neuronal cell bodies, is clearly distinguishable from the surrounding white matter (Fig. 5b). The gray matter is organized into the dorsal and ventral horns. The white matter is divided into regions such as the lateral funiculus (or column) and ventral funiculus. The central canal is located at the center of the spinal cord and is lined by ependymal cells (Fig. 5b).

### Sensory system

#### Eye

The eye of *G. rufa*, like that of other teleosts, is a laterally positioned, highly specialized sense organ^54,55^. The eyes are composed of the cornea, lens, iris, retina, retinal pigment epithelium, choroid, and sclera (Fig. 5c). Reflecting the adaptation for aquatic vision, the eye of *G. rufa* has a spherical, highly refractive lens (Fig. 5c) as the primary refractive structure, similar to other teleosts^38^. The retina is a sheet of neural tissue lining the posterior part of the eye (Fig. 5c). The retina consists of organized neural layers and underlain by the choroid, which supplies nutrients and absorbs excess light. Overall, the ocular structure of *G. rufa* is closely similar to that of zebrafish (Fig. S5s).

#### Olfactory sac

The primary function of the teleost olfactory sac is to detect molecules in the aquatic environment^56^. This function plays a critical role in feeding, predator avoidance, and mating behavior^57^. The olfactory sac is located on the dorsal side of the head (Fig. S4a). It consists of a pair of cavities that communicate with the outside through the external nostrils (Fig. S4e). To increase surface area, lamellae protrude into the olfactory sac (Fig. 5d). The olfactory sac is lined mainly by ciliated columnar epithelial cells (olfactory cells) (Fig. 5d). In the lamina propria, olfactory nerve fibers originating from the olfactory epithelium gather into fascicles (Fig. 5d).

#### Inner ear

The inner ear functions in hearing and maintaining balance^58,59^. In the auditory system of *G. rufa*, neither an outer ear nor a middle ear is present (Fig. S4d), as in other teleosts^28^. In the otolith organs, the otoliths, which are calcified stones, lie on the surface of the sensory epithelium (Fig. 5e). The sensory epithelium consists of the ciliated hair cells. The hair cells are activated when the otolith is displaced by gravitational forces or low-frequency vibrational stimuli^58,59^.

#### Taste buds

The taste buds detect chemical stimuli in the aquatic environment, facilitating food recognition and selection^60^. In *G. rufa*, taste buds are found in the oropharyngeal epithelium and on the surface of the lip. The sensory cells are elongated and exhibit apical microvilli projecting into the lumen (Fig. 5f). In zebrafish, taste buds are also distributed in the epithelium of the lip and the oropharyngeal epithelium, and microscopically the shape of the taste buds is similar to that of *G. rufa* (Fig. S5v).

#### Lateral line

The lateral line is a mechanosensory organ in the skin that detects water flow and vibration^61^. The lateral line canals run bilaterally along the trunk (Fig. 1b). Within the lateral line canals, there are neuromast cells, which are specialized mechano-receptive cells (Fig. 5g). The neuromast cells are specialized hair cells possessing sensory cilia that extend into the lumen (Fig. 5g). Water movement or vibration displaces these cilia, leading to activation of the hair cells and transmission of signals to the central nervous system^38^.

### Musculoskeletal system and skin

#### Skin

The skin forms the outer protective barrier covering the entire body of *G. rufa*. (Fig. 1b, 5h). The skin is composed of the epidermis and dermis. The epidermis of *G. rufa* is a stratified squamous epithelium. Keratinization of epidermal cells is rare except in the lip region (Fig. 2c). In the epidermis, *G. rufa* has a higher abundance of goblet cells compared to zebrafish (Fig. 5i,S5w). Scales are embedded within the dermis (Fig. 5i,S5w).

#### Skeletal muscle

The bulk of *G. rufa* body consists of segmented skeletal muscles (Fig. 1d,S2,S4). Skeletal muscle consists of long, thin, fibrous myofibers, which are multinucleated cells (Fig. 5j). The nuclei of myofibers are located at the periphery of the myofibers (Fig. 5j). Myofibers contain filamentous components in their cytoplasm known as myofibrils.

#### Bone and cartilage

Ossified tissue provides the structural framework of the body. Ossified tissue encases the brain for protection and forms the vertebral column along the body axis (Fig. 1c). The fins are also supported by ossified elements (Fig. 1c). Cartilage is observed in the gill filaments (Fig. 4a) and intervertebral regions (Fig. 5k).

### High-temperature resistance

To quantitatively elucidate how *G. rufa* and zebrafish, a widely used model teleost, respond differently to human body temperature (37 °C), we conducted two physiological experiments. First, we evaluated the time course of survival rates of *G. rufa* and zebrafish at 37°C. While *G. rufa* exhibited a modest decline in survival, with less than 20% mortality observed by day 20, the survival curve plateaued thereafter (Fig. 6a). In contrast, zebrafish showed a rapid decrease in survival. The difference in survival between the two species was statistically significant (n = 12, *p* < 0.0005, log-rank test).

**Figure 6.**
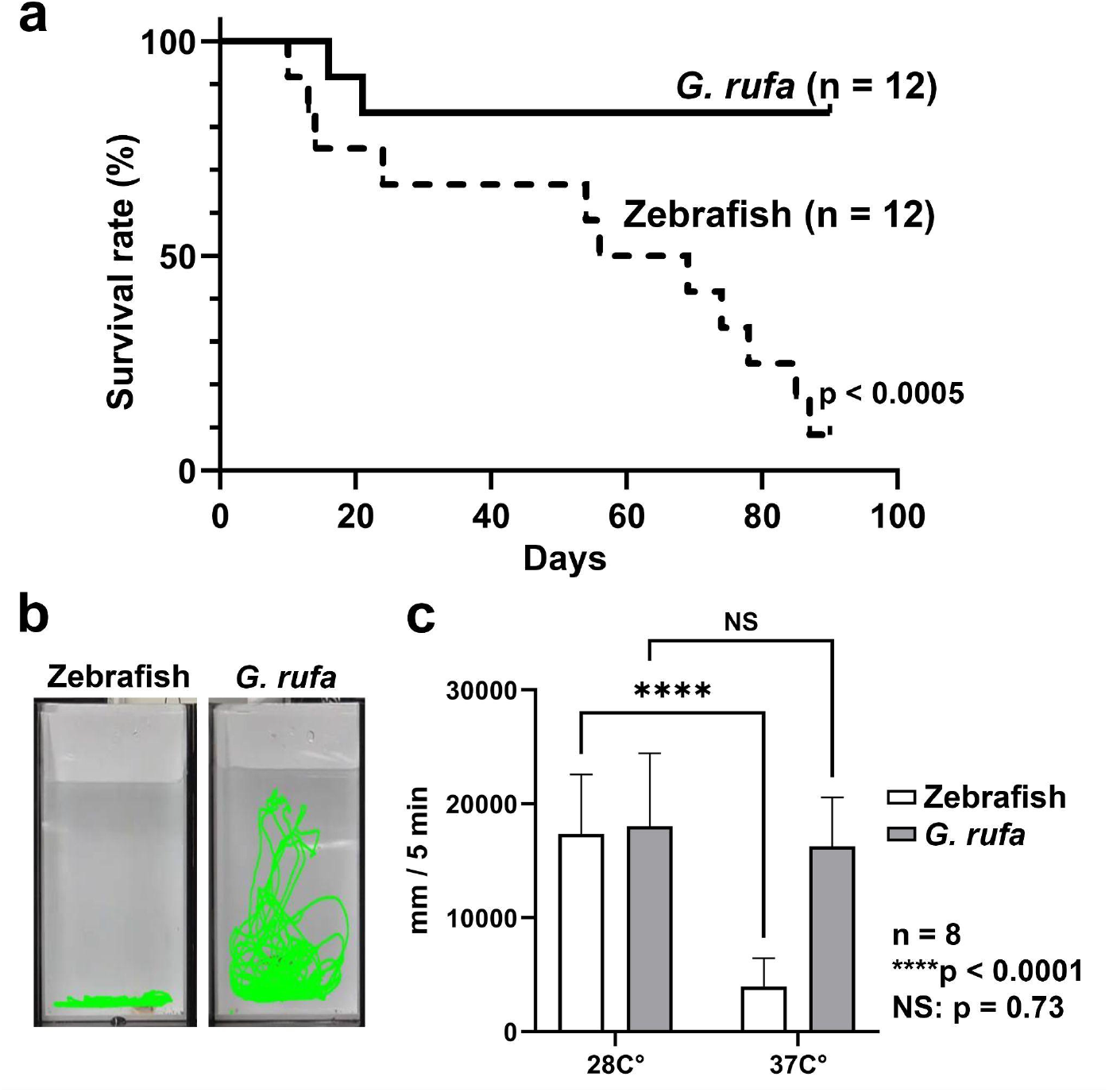
Responses of *G. rufa* and zebrafish to 37 °C. (**a**) Kaplan-Meier survival curves of *G. rufa* and zebrafish at 37 °C. While *G. rufa* maintained high survival over 90 days, zebrafish exhibited a rapid and progressive decline in survival (n = 12 for each group; log-rank test, *p* < 0.0005). (**b**) Swimming trajectories recorded for 5 min following feeding stimulation at 37 °C. Green lines indicate the swimming trajectories of individual fish. (**c**) Quantification of total swimming distance (mm/5 min) at 28 °C and 37 °C for each species. Data are presented as mean ± SD. n = 8; unpaired Student’s t-test; **** *p* < 0.0001; NS, not significant.

Next, we analyzed swimming distances following feeding stimulation (Fig. 6b). In *G. rufa*, there was no significant difference in swimming distance between the 28 °C and 37 °C conditions (Fig. 6c, n = 8, unpaired Student’s t-test, *p* = 0.73). In contrast, zebrafish exhibited a significant reduction in swimming distance at 37 °C compared to 28 °C (Fig. 6c, *n* = 8, unpaired Student’s t-test, *p* < 0.0001). These results indicate that zebrafish are considerably more vulnerable to human body temperature (37 °C) in terms of both survival and behavior, whereas *G. rufa* remains physiologically stable under the same conditions.

## Discussion

In this study, we performed comprehensive histo-anatomical analyses of the whole body of *G. rufa*, thereby providing an integrated description of its overall body organization. In addition, through comparison with zebrafish, a representative small teleost model organism, we characterized both morphological similarities and differences and quantitatively evaluated physiological responses to 37 °C, a temperature corresponding to the human body temperature. The morphological and physiological insights obtained in this study, together with the recently reported chromosome-level genome assembly^23^, are expected to serve as foundational resources for establishing this species as a new biomedical model. Furthermore, the morphological differences revealed in this study, such as intestinal length, combined with the difference in thermal tolerance, offer valuable comparative insights into environmental adaptation and evolutionary diversification within cyprinid teleosts.

Over the past two decades, small teleosts such as zebrafish and medaka have been widely used as human disease models. These small fish models offer significant advantages for high-throughput *in vivo* analyses and drug screening, not only because they are subject to fewer ethical constraints than mammalian or primate models, but also because their basic anatomical and physiological features are broadly conserved across vertebrates, including humans^7,62,63^. Indeed, successful translational applications have been reported, including the discovery that prostaglandin E2 regulates hematopoietic stem cell homeostasis^64^, which led to clinical trials for cord blood transplantation^65^, and the identification of novel therapeutic candidates for Dravet syndrome using a zebrafish model^66^. Furthermore, the efficacy of existing drugs, such as ezetimibe, has been re-evaluated using a medaka model of non-alcoholic steatohepatitis^67^, with some findings advancing to clinical application.

The comprehensive histological analyses in this study, combined with the results of the preceding genome sequencing project^23^, demonstrate that *G. rufa* possesses genomic and organ structures highly similar to those of zebrafish. This strongly suggests that the vast accumulation of research findings and experimental methodologies established in small fish models can be directly reproduced and applied to *G. rufa*. Moreover, *G. rufa* holds the potential to surpass existing models due to its unique amenability to experimentation at 37 °C. Since human biological processes are extensively optimized for 37 °C, experiments conducted in lower-temperature environments possess inherent limitations. First, in cancer research, key processes such as the proliferation, metastasis, and angiogenesis of human tumor cells are temperature-dependent; thus, modeling these processes at lower temperatures carries the risk of underestimating tumor aggressiveness. Second, regarding infectious diseases, many human pathogens are adapted to proliferate at body temperature. Furthermore, the host immune response, particularly immune activation during fever, is physiologically promoted at temperatures of 37–40 °C. Consequently, ectothermic models maintained at 28 °C cannot fully capture the dynamic interaction between pathogen and host. Finally, in metabolism and pharmacology, basic enzymatic reactions follow the Q10 temperature coefficient rule, meaning a temperature difference of approximately 9 °C (37 vs 28 °C) drastically alters metabolic rates. Therefore, in drug discovery, it is essential to evaluate pharmacokinetics under conditions that mimic human thermal physiology. From these perspectives, *G. rufa* is expected to serve as a novel model organism that overcomes the limitations of existing small fish models, enabling analyses under physiological conditions that more closely resemble those of humans.

Another notable feature identified in this study is the markedly elongated intestine of *G. rufa*. In contrast to the relatively simple intestinal morphology of zebrafish^68^, elongated intestinal structures have also been reported in other species of the genus *Garra*^*29*^, suggesting that this trait may be conserved within the genus. This characteristic is likely related to the benthic feeding ecology of *G. rufa*, which consumes algae and biofilms in substrate-associated environments^69^. An extended intestine may provide sufficient residence time for the efficient digestion of such dietary components. While the zebrafish intestine is known to be subdivided into anterior, mid, and posterior segments, it remains unclear whether a comparable segmentation occurs during the development of *G. rufa* or whether intestinal elongation proceeds through a distinct developmental program. Investigating the intestinal development of *G. rufa* will be essential to clarify how the elongated intestine is formed and to elucidate the underlying developmental and molecular mechanisms.

While this study established the superior thermal tolerance of *G. rufa*, the underlying molecular mechanisms remain to be elucidated. The absence of distinguishing morphological features accounting for such thermal tolerance suggests that the thermal tolerance of *G. rufa* relies on intrinsic cellular properties, such as stress-responsive pathways, rather than gross anatomical properties. Given the high genomic conservation between the two species^23^, this difference likely arises from divergence in gene regulatory mechanisms. Future comparative analyses of gene regulatory networks and functional validation are therefore warranted to identify the specific genetic drivers of this adaptation.

To further develop *G. rufa* as an experimental model organism, the establishment of laboratory-maintainable strains with uniform genetic backgrounds suitable for molecular biological analyses will be essential. In addition, the development of reliable breeding and egg collection systems, analogous to those established for zebrafish^70^, as well as the implementation of genetic manipulation techniques such as transgenesis and CRISPR/Cas9-mediated genome editing, will be critical. Moreover, extending the imaging capabilities beyond the micro-CT and MRI approaches used in this study to include tissue clearing for whole-body single-cell imaging^71,72^ and spatial transcriptomics^73^ would greatly enhance high-resolution spatial phenotyping. By establishing these experimental tools in *G. rufa* and fully utilizing the recently released chromosome-level genome sequence^23^, *G. rufa* has the potential to emerge as a complementary biomedical model that enhances the experimental utility of zebrafish.

## Methods

### Fish maintenance and sample collection

Adult *G. rufa* and zebrafish (AB strain) were maintained in laboratory aquaria at 28 °C under a 14:10 h light–dark cycle. *G. rufa* were fed once daily with Hikari Crest Pleco (Kyorin, Himeji, Japan), while zebrafish were fed twice daily with Gemma Micro 300 (Skretting, Stavanger, Norway). All animal handling and experimental procedures were conducted in accordance with institutional guidelines approved by the Animal Care and Use Committee of the Mie University (approval no. 2024-57). For histological and imaging analyses, healthy adult fish were anesthetized with 0.1% 2-phenoxyethanol (Fujifilm Wako Pure Chemicals, Tokyo, Japan) and euthanized by rapid cooling on ice.

### Gross anatomical observation and imaging

External morphology was observed using an SMZ745T stereomicroscope (Nikon, Tokyo, Japan) equipped with a TrueChrome II Plus Full HD HDMI camera (Biotools, Gunma, Japan). For macro imaging of large specimens, including whole-body views, a Tough TG-6 digital camera (Olympus, Tokyo, Japan) was employed in macro mode.

### Micro-computed tomography (µ-CT)

For three-dimensional skeletal visualization, whole-body µ-CT scanning was conducted using a Cosmo Scan CT AX system (Rigaku, Tokyo, Japan). The scanning parameters were set as follows: tube voltage, 90 kV; tube current, 88 µA; and exposure time, 4 min. The resulting images were reconstructed with an isotropic voxel size of 90 µm using Cosmo Scan CT Viewer software (Rigaku).

### Magnetic resonance imaging (MRI)

MRI was performed using a small animal MRI system (MRVivo LVA Academic Model 1506; Aspect Imaging, Shoham, Israel) equipped with a 30 mm diameter RF coil. Samples were placed on a custom-made stage consisting of a 50 mL centrifuge tube filled to 60–70% capacity with solidified 1% agarose. Images were acquired using a 2D Fast Spin Echo sequence with a flip angle of 90° and a matrix size of 128 × 256. Image reconstruction and analysis were carried out using ImageJ software (Fiji version 1.54p; NIH, Bethesda, MD, USA).

### Comparison of intestinal length between *G. rufa* and zebrafish

To quantify normalized intestinal length, we first measured the standard body length^74^, calculated as the total body length minus the caudal fin length. The entire intestine was then surgically excised, gently straightened without stretching, and measured from the anterior to posterior end to obtain the total intestinal length. The normalized intestinal length was calculated by dividing the intestinal length by the standard body length. Using this procedure, we measured the normalized intestinal length in three adult *G. rufa* and three adult zebrafish. Statistical analyses were performed in R v4.1.2. Normality of each group was tested using the shapiro.test function, and equality of variances between the two species was tested using the var.test function. As no significant deviation from normality or variance inequality was detected, a Student’s t-test assuming equal variances was conducted to compare the two species using the t.test function with the option “var.equal = TRUE”. P-values less than 0.05 were considered statistically significant.

### Histological preparation and staining

Whole fish or dissected tissues were fixed overnight in 4% paraformaldehyde in phosphate-buffered saline (4%PFA; Falma, Tokyo, Japan) at 4 °C. The fixed samples were sent to the Division of Medical Research Support, Advanced Research Support Center (ADRES), Ehime University, where they were processed for paraffin embedding, sectioning (5 μm thickness), and routine hematoxylin and eosin (H&E) staining. Stained sections were dehydrated, mounted with coverslips, and imaged in brightfield mode using a BZ-X710 fluorescence microscope (Keyence, Osaka, Japan).

### Peripheral blood smear and staining

Peripheral blood was collected from the dorsal aorta of anesthetized adult *G. rufa* using a heparinized glass capillary, following a method previously described for zebrafish ^75,76^. Blood smears were prepared immediately on clean glass slides, air-dried, and stained with Wright-Giemsa Stain (ScyTek Laboratories, Logan, UT, USA) for cytological evaluation. Stained slides were examined under a BX51 light microscope (Olympus, Tokyo, Japan) equipped with an HD Lite 1080P digital camera (Tucsen Photonics, Fuzhou, China).

### Zebrafish reference histological images

Histological reference images used for comparison were obtained from the zebrafish virtual slides of the Bio-Atlas (https://bio-atlas.psu.edu/). Specific frames were cited using the “link to this” tool provided on the website. We acknowledge the Bio-Atlas project (NIH grant 5R24 RR01744), the Jake Gittlen Cancer Research Foundation, and the Pennsylvania Tobacco Settlement Fund as the sources of these materials^26^.

### Survival assay

For the evaluation of the thermal tolerance, *G. rufa* and zebrafish individuals were exposed to 37 °C. In each group, 12 individuals per species were used for the survival assay. The individuals were collectively maintained in a 10 L tank under a controlled condition for each group. Survival curves were generated using the Kaplan–Meier method and compared by the log-rank (Mantel-Cox) test using Prism version 10.6.1 (GraphPad Software, Boston, MA, USA).

### Swimming activity assay

After housing for 2 weeks at 28 °C or 37 °C, swimming behavior in response to feeding stimulation was evaluated. Individual *G. rufa* and zebrafish were transferred into rectangular tanks containing water equilibrated to the respective temperature (28 °C or 37 °C). Following a 10-min acclimation period, fish were fed by Gemma Micro 300 (Skretting), and swimming activity was recorded for 5 min. Videos were captured at 30 frames per second (fps) using a high-definition digital video camera (HC-V495M; Panasonic, Osaka, Japan) positioned in front of the tank. Swimming trajectories were analyzed using ToxTrac software^77^ to quantify total swimming distance (mm/5 min). Statistical comparisons between temperatures within each species were performed using an unpaired Student’s t-test in Prism version 10.6.1 (GraphPad Software). A *p*-value less than 0.05 was regarded as statistically significant.

## Acknowledgments

The authors are grateful to Dr. Natsuki Morimoto at the Fisheries Technology Institute, Japan, for her valuable histological advice. The authors also thank Dr. Kiki Syaputri Handayani, Dr. Victor David Nico Gultom, and Dr. Fahrurrozi at the Research Center for Marine and Land Bioindustry, Research Organization for Earth Science and Maritime (BRIN) for their advice on *G. rufa* husbandry. The authors thank the Division of Medical Research Support, Advanced Research Support Center (ADRES), Ehime University, for their technical assistance with histological analysis, and Ms. Masami Yamamura at Mie University for her assistance with *G. rufa* breeding. Part of this work was supported by “Advanced Research Infrastructure for Materials and Nanotechnology in Japan (ARIM)” of the Ministry of Education, Culture, Sports, Science and Technology (MEXT) (Proposal Number JPMXP1225NU0424).

## Author Contributions

YS and TK conceived and designed the project. YS, LZ, and SN performed the experiments. TK, KK, SN, LZ, and YS performed the histo-anatomical analyses. TK, KK, SN, LZ, and YS wrote the manuscript. TK, KK, OS, and YS reviewed the overall manuscript structure and interpretation. All authors read and approved the final version of the manuscript.

