## Supplementary figures for "Histo-anatomical atlas and thermal tolerance of *Garra rufa*: A novel small teleost model adaptable to human body temperature"

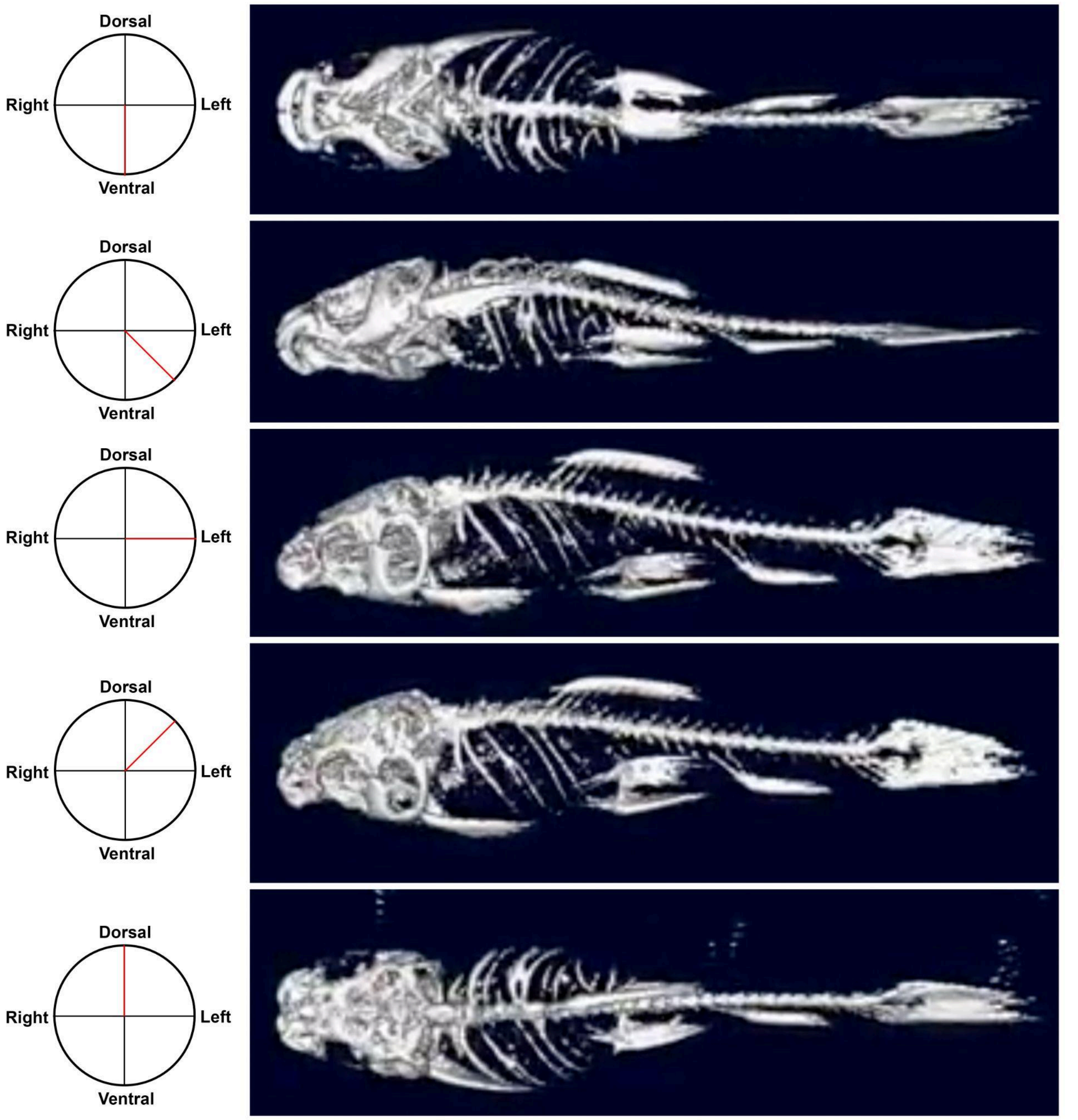


**Figure S1. Whole-body skeletal μCT images**

Representative μCT images showing the whole-body skeletal structure from different rotational views. The red lines in the circular diagrams indicate the orientation of each view relative to the dorsal–ventral and left–right axes.

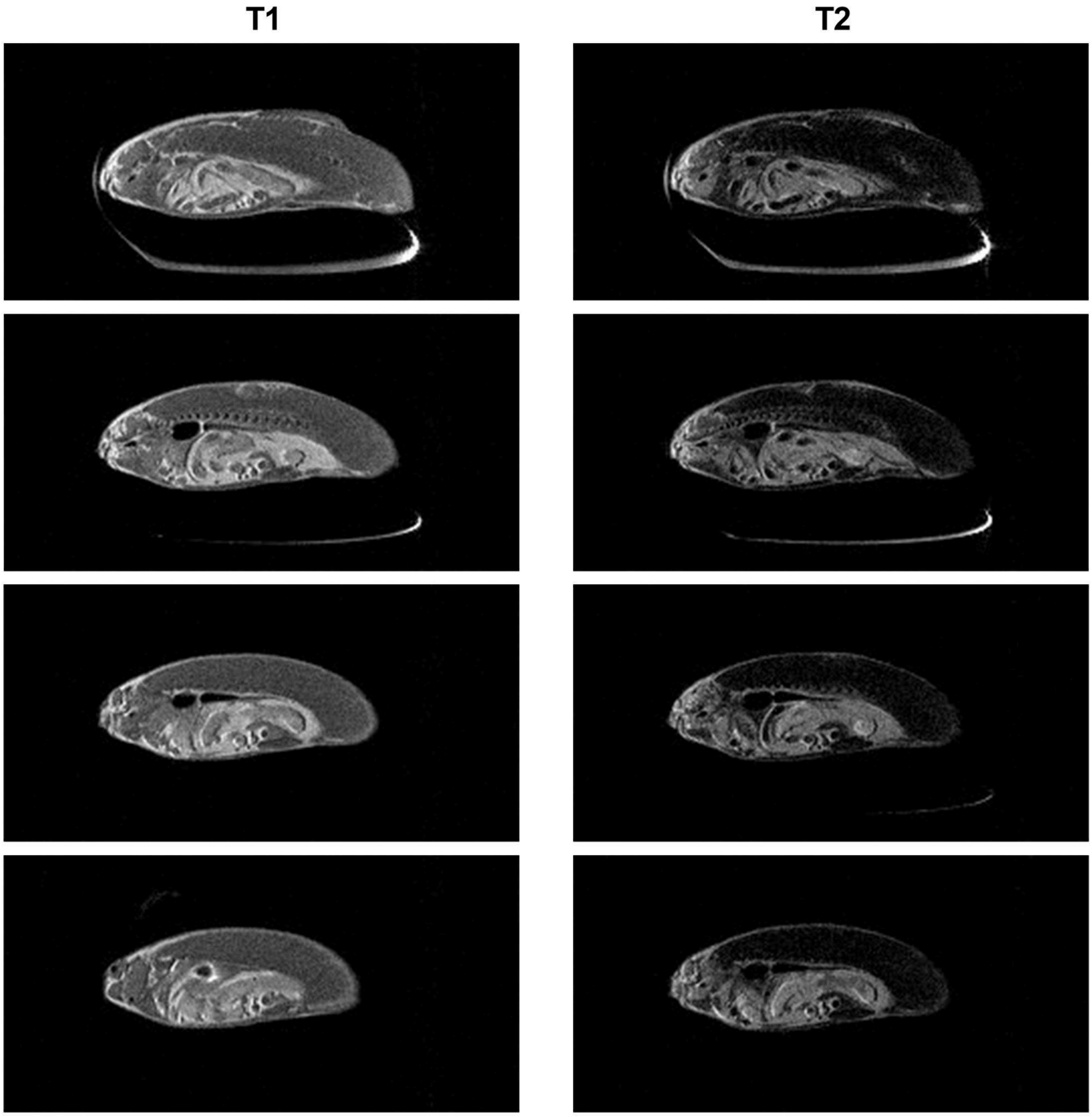


**Figure S2. Whole-body skeletal MRI images**
The left column shows T1-weighted images, and the right column shows T2-weighted images.

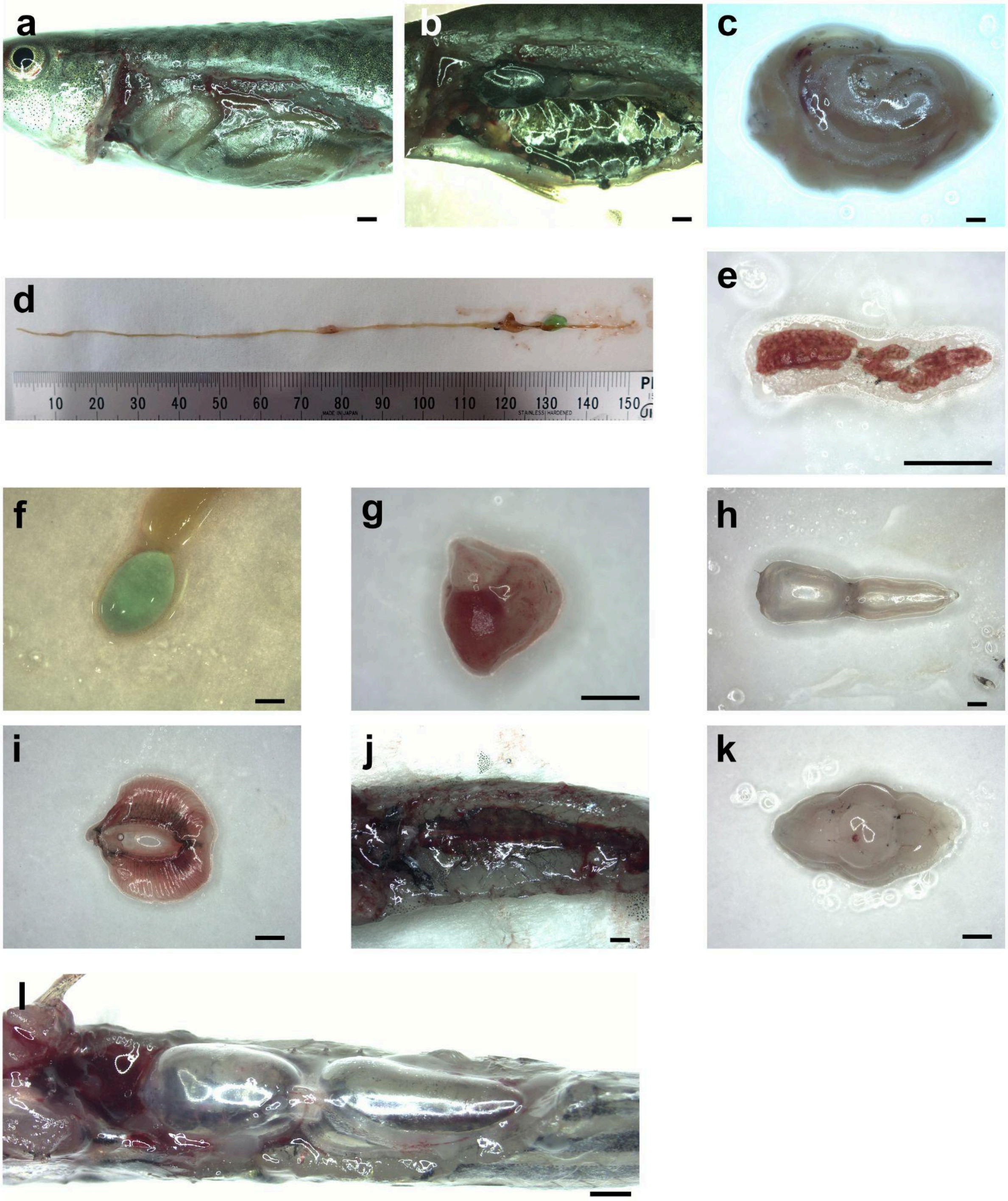


**Figure S3. Whole-body gross anatomical findings in *G. rufa***

(a) Intraperitoneal organs after abdominal wall resection of *G. rufa*. (b) Peritoneal cavity and its lining peritoneum of *G. rufa*. (c) Liver and intestine of *G. rufa*. Adipose deposition is observed in the mesentery. (d) Intestine of *G. rufa*. (e) Spleen of *G. rufa*. (f) Gallbladder of *G. rufa*. (g) Heart of *G. rufa*. (h) Swim bladder of *G. rufa*. (i) Gills of *G. rufa*. (j) Kidney of *G. rufa*. (k) Brain of *G. rufa*. (l) Abdominal cavity of Zebrafish. Scale bars represent 1 mm.

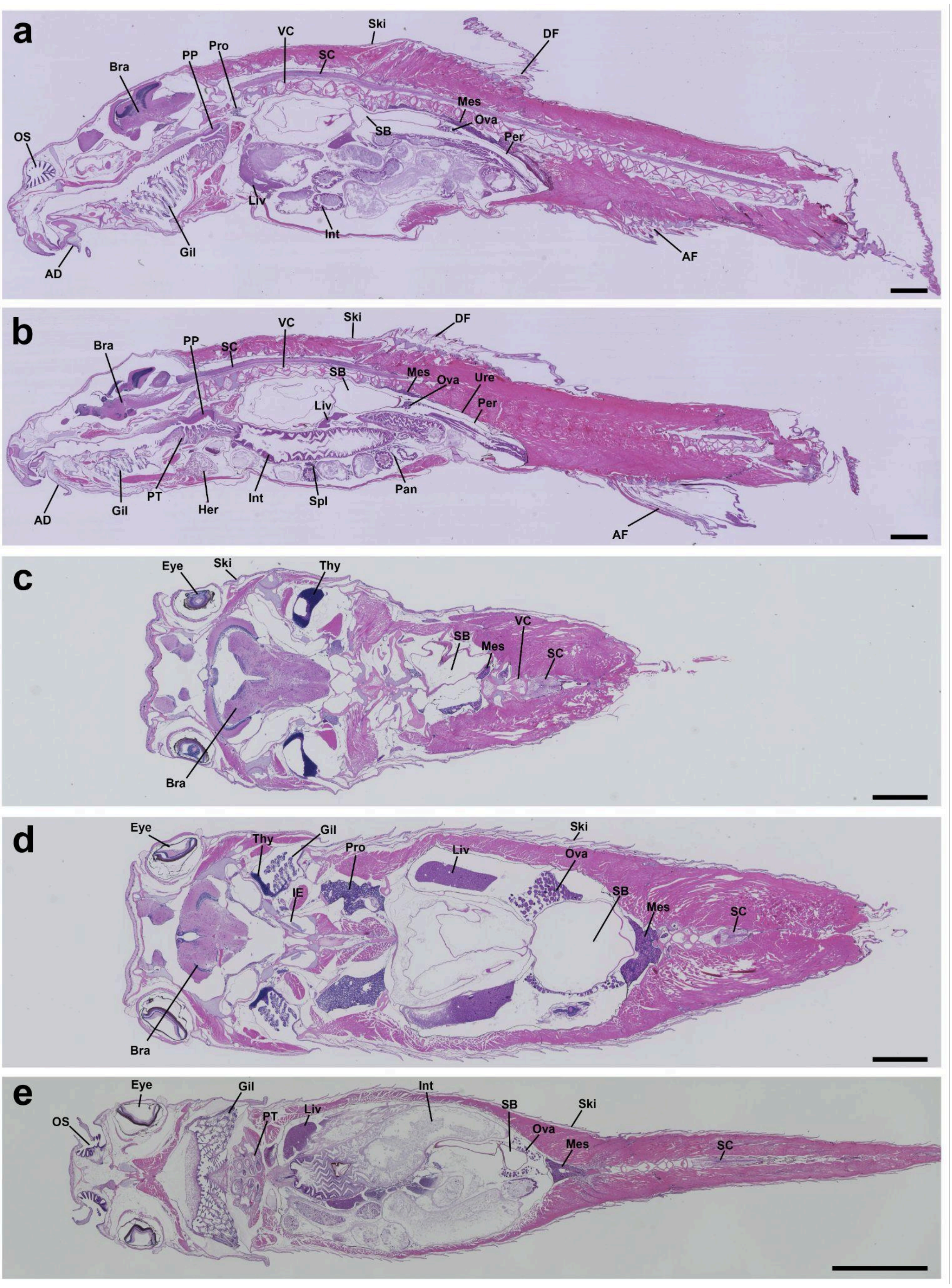


**Figure S4. Representative whole-body sagittal sections**

Abbreviations: Adhesive disc, AD; AF, anal fin; Bra, brain; DF, dorsal fin; Gil, gills; Her, heart; IE, inner ear; Int, intestine; Liv, liver; Mes, mesonephros; OS, olfactory sac; Ova, ovary; Pan, pancreas; Per, peritoneum; PP, pharyngeal pad; Pro, pronephros; PT, pharyngeal tooth; SC, spinal cord; Spl, spleen; SB, swim bladder; Ski, skin; Thy, thymus; Ure, ureter; VC, vertebral column. Scale bars represent 1 mm.

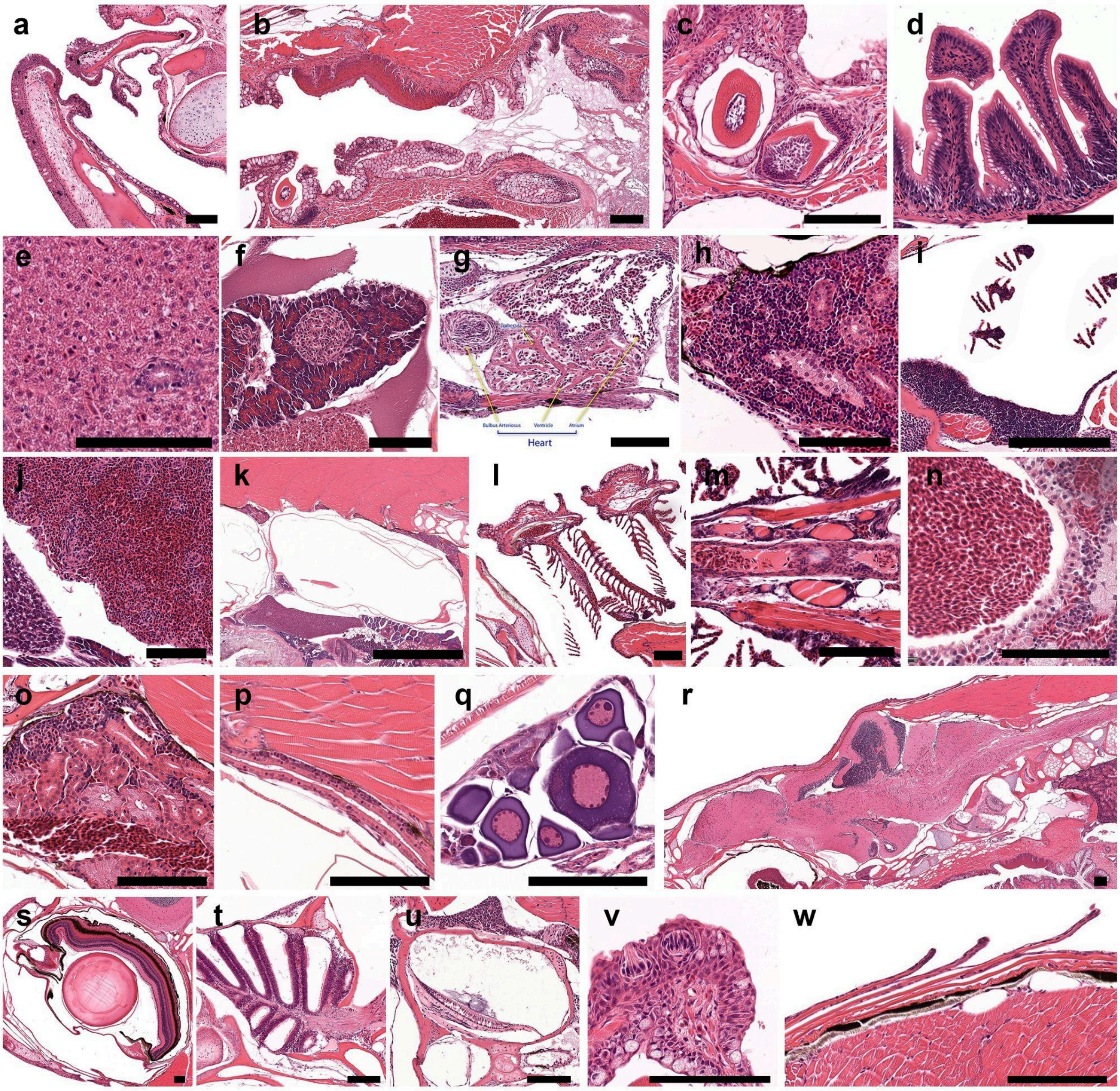


**Figure S5. Reference Histological Sections of Zebrafish Retrieved from the Bio-Atlas**
Zebrafish reference histological images were retrieved from the zebrafish virtual slides of the Bio-Atlas (https://bio-atlas.psu.edu/). For details regarding data source and attribution, refer to the Materials and Methods section. (a) Mouth. (b) Pharynx and esophagus. (c) Pharyngeal tooth. (d) Intestine. (e) Liver. (f) Pancreas. (g) Heart. (h) Pronephros. (i) Thymus. (j) Spleen. (k) Swim bladder. (l) Gills. (m) Thyroid. (n) Adrenal tissue. (o) Mesonephros. (p) Ureter. (q) Ovary. (r) Brain and spinal cord. (s) Eye. (t) Olfactory sac. (u) Inner ear. (v) Taste buds in the pharyngeal epithelium. (w) Skin. Scale bars represent 100 μm.
